# Genomic analysis reveals the presence of emerging pathogenic *Klebsiella* lineages aboard the International Space Station

**DOI:** 10.1101/2023.05.05.539530

**Authors:** Georgios Miliotis, Nitin Kumar Singh, Francesca McDonagh, Louise O’Connor, Alma Tuohy, Dearbháile Morris, Kasthuri Venkateswaran

## Abstract

*Klebsiella* species, including *Klebsiella pneumoniae, Klebsiella aerogenes*, and *Klebsiella quasipneumoniae*, are opportunistic pathogens that are known to cause infections in humans. Hypervirulent *Klebsiella pneumoniae* (hvKP) is a subgroup of *K. pneumoniae* that has gained attention due to its global dissemination and its ability to cause invasive infections in community settings amongst immunocompetent individuals as well as its increasing levels of antibiotic resistance. Our study reports the first complete genotypic analysis including mobile genetic elements (MGEs) of *Klebsiella* isolates from the International Space Station (ISS). The genomes of *K. pneumoniae, K. aerogenes*, and *K. quasipneumoniae* provided valuable insights into their antimicrobial resistance, virulence, thermotolerance, disinfectant resistance, and MGEs. All isolates belonged to emerging lineages with pathogenic potential, with *K. quasipneumoniae* ST138 presenting spatial and temporal persistence aboard the ISS, possibly due to its genotypic profile encoding for numerous resistance genes to disinfectants and heavy metals. We also report on the isolation of a yersiniabactin encoding *K. pneumoniae*, belonging to the emerging high-risk ST101 clone, aboard the ISS. Potential dissemination of hvKp strains on ISS could pose a putative risk to the immunocompromised crew. The presence of MGEs containing virulent loci could facilitate horizontal gene transfer to other benign microorganisms on the ISS, potentially increasing their virulence. In addition, genetic divergence from their respective lineages for some *Klebsiella* genomes was predicted and hypothesized to be due to the unique spaceflight environmental pressures. These findings highlight the importance of monitoring problematic microbial communities in space to understand their surviving abilities and potential impact on human health.

**Importance:** The International Space Station (ISS) is a unique hermetically sealed environment that poses environmental pressures not encountered on Earth, including microgravity and radiation While the adaptability of bacteria during spaceflight is not fully understood, recent research has suggested that it may be species and even clone specific. Given the spaceflight-induced suppression of the human immune system, it is essential to understand the genomics of potential human pathogens in spaceflight. Such understanding could provide valuable insights into species and lineages of medical astromicrobiological importance. Here, we used hybrid assembly approaches and comparative genomics to provide the first comprehensive genomic characterisation of 10 *Klebsiella* isolates retrieved from the ISS. Our findings revealed that *K. quasipneumoniae* ST138 exhibits spatial and temporal persistence aboard the ISS, with evidence of genomic divergence from the ST138 lineage on Earth. Additionally, we characterized plasmids from *Klebsiella* species of ISS origin, which encoded disinfectant and thermoresistance genes suggesting that these might aid adaptability. Furthermore, we identified an MGE containing a hypervirulence-associated locus belonging to a *Klebsiella pneumoniae* isolate of the “high risk” ST101 clone. Our work provides valuable insights into the adaptability and persistence of *Klebsiella* species during spaceflight, highlighting the importance of understanding the behaviour of potential pathogenic bacteria in space.

## 1. Introduction

The genus *Klebsiella* includes eight species of facultative anaerobic Gram-negative bacteria that are ubiquitously present in the natural environment(1). They can also be found as human and animal gastrointestinal commensals. Some species of the genus *Klebsiella* are well-established human opportunistic pathogens known to cause both community and hospital-acquired infections (HAIs)(2, 3).

*Klebsiella pneumoniae* is clinically the most important *Klebsiella* species, having a multifaceted pathogenicity profile, including urinary tract infections (UTIs), meningitis, pneumonia, and sepsis(4). *K. pneumoniae* is of concern due to multidrug resistance (MDR), extended drug resistance (XDR), and hypervirulence (hvKp) associated lineages causing outbreaks in community and healthcare settings(5, 6). The dispersal of strains belonging to hvKp lineages is of an increasing concern to the medical community(7). HvKp differs from classic *K. pneumoniae* (cKp) due to its ability to infect immunocompetent individuals in community settings. Infections associated with hvKp commonly affect multiple sites due to metastatic spread(8). Although there is not a clear definition for hvKp, there are several siderophore virulence factors reported to be associated with a hvKp pathotype, mainly; aerobactin (iuc), salmochelin (iro) and yersiniabactin (ybt) (9, 10). Recently, Kleborate, a genotyping tool has been developed to assess and quantify virulence amongst the *K. pneumoniae* species complex using a 5-tier scoring system based on the presence of key virulence associated loci (10). Overall, these virulence loci are commonly identified on mobile genetic elements (MGEs) such as self-mobilizable plasmids and integrative and conjugative elements (ICEs). Among *K. pneumoniae* populations, ICEKp is known to mobilise the *ybt* locus which encodes for the key iron scavenging virulence factor yersiniabactin along with its receptor(11). Besides virulence factors, capsular serotypes K1 and K2 are more frequently encountered in hvKp(12). Other capsular types, such as KL106, are associated with globally disseminated pathogenic *K. pneumoniae* strains, but their structures have not been fully resolved (13).

Other *Klebsiella* species are also known to cause HAIs but rarely infect immunocompetent individuals outside of healthcare associated settings. *Klebsiella aerogenes* is often associated with HAIs in immunocompromised patients and infections primarily include pneumonia, osteomyelitis, endocarditis, and UTIs (14) *K. aerogenes* is known to produce a chromosomal cephasolosporinase (AmpC) exhibiting resistance to penicillins as well as 1^st^ and 2^nd^ generation cephalosporins(15). AmpC β-lactamases are not inhibited by β-lactamase inhibitors such as clavulanic acid, and as such, β-lactamase inhibitor combinations are not recommended for the treatment of *K. aerogenes* associated infections(16).

*Klebsiella quasipneumoniae* is separated into two subspecies *K. quasipneumoniae* subsp. *quasipneumoniae* and *K. quasipneumoniae* subsp. *similipneumoniae*. Both subspecies have now been recognized as human opportunistic pathogens(17). Recent reports have identified *K. quasipneumoniae* subsp. *similipneumoniae* as a causative agent of neonatal septicaemia outbreaks in China and Nigeria(18, 19). An increased awareness of its dissemination and pathogenicity is therefore required.

The International Space Station (ISS) is a hermetically sealed extreme environment which puts microorganisms under the unique selective pressures of microgravity and radiation (ionising/UV). During the Microbial Tracking-1 experiment, which was dedicated to characterizing the ISS microbiome, 10 strains belonging to three different *Klebsiella* species were identified(20). Using culture-independent methods, it has been reported that *K. pneumoniae* can be identified in multiple locations on the ISS, and its succession over time has been documented (21, 22). The overall importance of understanding the genomic background, evolution and persistence of potentially pathogenic microorganisms during spaceflight, in terms of medical astromicrobiology has also been recently discussed(23).

The aim of this research communication is twofold: first, to generate complete genomes and circular chromosomal maps of the ten *Klebsiella* strains isolated from the ISS. These genomes would be used to investigate the strains’ antimicrobial resistance, virulence, thermotolerance, heavy metal resistance, and mobile genetic elements (MGEs), and would provide the first complete *Klebsiella* plasmids from the ISS. Second, large-scale population genomics would be employed to understand the evolution and potential divergence of the ISS *Klebsiella* isolates from their respective lineages.

## 2. Results

### 2.1. Antimicrobial susceptibility testing

Most clinically relevant antibiotics were found to be effective against all tested *Klebsiella* strains. As expected, the strains exhibited resistance to ampicillin and rifampicin. Among the strains, three displayed resistance to second-generation cephalosporins, while five exhibited intermediate resistance to quinolones, two to aminoglycosides, and one to the carbapenem imipenem. (**Table S1**).

### 2.2 Assembly statistics

Due to hybrid assembly using Illumina and ONT sequencing platforms, circular plasmid and closed chromosomal genomes were possible. The assembly statistics are provided (**Table S2**). A direct genomic comparison of the chromosomes of all 10 ISS *Klebsiella* strains is shown in (**Fig.1A**).

**Figure 1:**
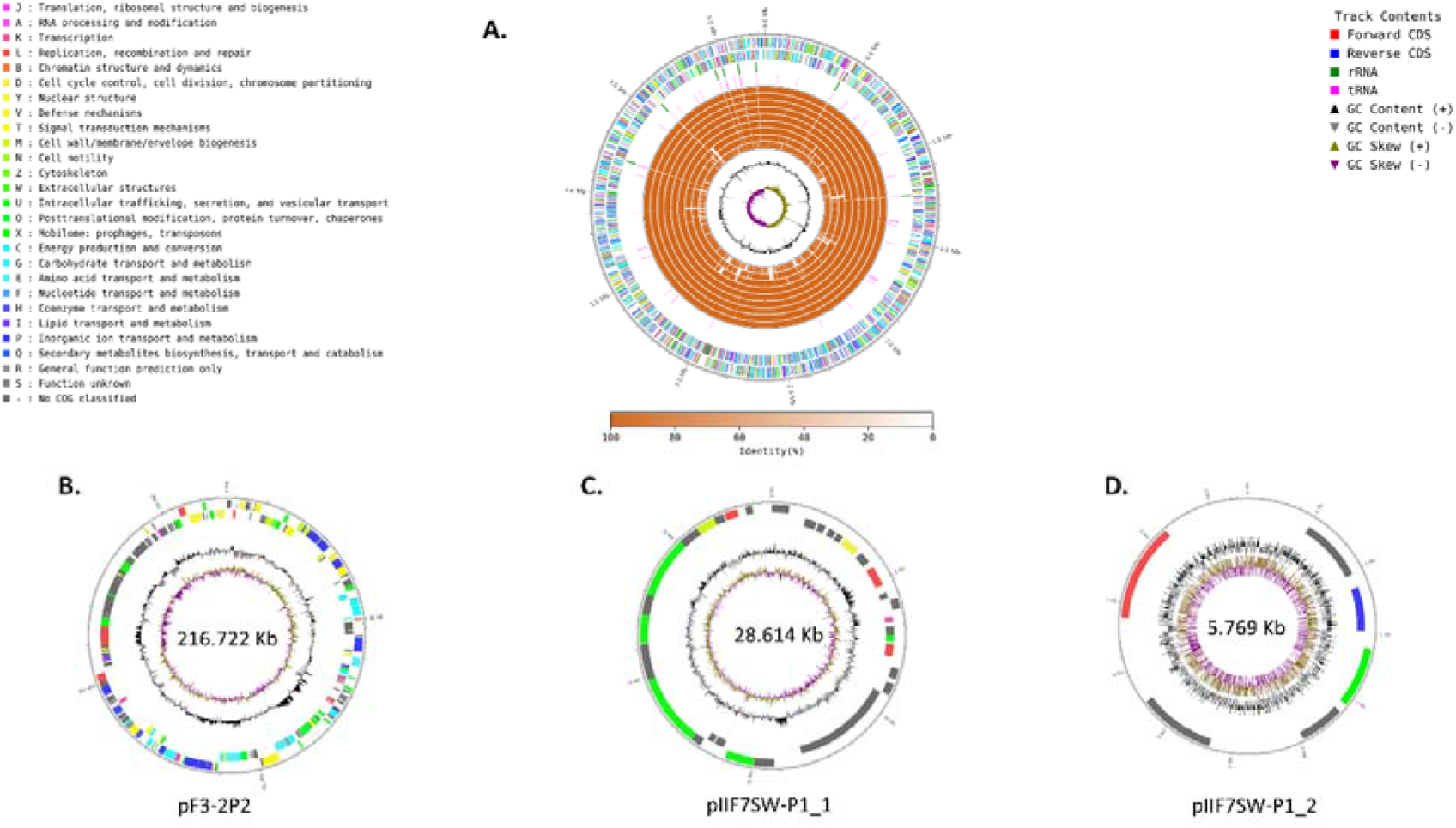
Genetic comparison and functional characteristics of *Klebsiella* genomes and plasmids. **A:** Direct genomic comparison for all (n=10) *Klebsiella* genomes retrieved from the ISS. The two outermost circles represent the forward and reverse strands of *K. quasipneumoniae* strain IF1SW-B2. Each gene is coloured per COG function. The two next rings indicate rRNA and tRNA genes respectively. The inner circles represent regions of genomic similarity between IF1SW-B2 and the other ISS strains as follows: 1. F3-6P(2), 2. IF2SW-B3, 3. IF1SW-B2, 4.F3-6P(1), 5. IF2SW-P1, 6. IF1SW-P4, 7. IF1SW-P3, 8. IIIF3SW-P1, 9. F3-2P(2), 10. IIIF7SW-P1. The innermost circles represent GC content and GC skew respectively. **B.** Represents the 216.722 Kb plasmid pF3-2P(2) retrieved from *K. pneumoniae* strain F3-2P2(2). **C.** Represents the 28.614 Kb plasmid pIIIF7SW-P1_1 retrieved from *K. aerogenes* strain IIIF7SW-P1. **D.** Represents the 5.769 Kb plasmid pIIF7SW-P1_2 retrieved from *K. aerogenes* strain IIIF7SW-P1. Coding sequences are coloured per COG function.

### 2.3. Genome typing

Genotyping based sequence type (ST) characterisation of the strains identified four different STs. The most common ST amongst the *K. quasipneumoniae* subsp. *similipneumoniae* was ST138 (7/8 *K. quasipneumoniae* strains). These strains also shared identical K and O locus alleles (K48/O5). Strain IIIF3SW-P1 was typed as ST3234 with K16 and O3 surface antigens. The *K. pneumoniae* strain F3-2P(2) belongs to the ST101 lineage with KL106 and O1/O2v2, while *K. aerogenes* strain IIIF7SW-P1 to the ST103 lineage with KL119 and O3/O3a (**Table S3**).

### 2.4. Plasmid characterisation

*K. pneumoniae* strain F3-2P(2) was found to contain a 216.722Kb, IncFIIK (FIIK-9 allele) conjugative plasmid encoding for 234 coding sequences (CDSs; **Fig. 1B**). The plasmid was further characterized to reveal the presence of gene clusters predominantly associated with resistance to heavy metals and high temperatures. Genes responsible for mercuric resistance (*merA*), arsenical resistance (*arsA/B/C*), copper resistance (*copA/B/D/R*), copper and silver export systems (*cusA/B/C/F/R/S*), and cell growth at high temperatures (*cplC, htpX*) were present.

*K. aerogenes* strain IIIF7SW-P1 was identified to carry two previously uncharacterised plasmids: pIIF7SW-P1_1 is a 28.614Kb, low copy (0.68 copies per cell), mobilizable plasmid identified to contain 38 CDSs (**Fig. 1C**). pIIF7SW-P1_2 is a 5.796Kb, high copy (16.10 copies per cell), non-mobilizable plasmid contained 6 CDSs (**Fig. 1D**; **Table S4**). pIIF7SW-P1_2 was found to encode for the heat shock gene *htrC*.

### 2.5. Average nucleotide identity analysis of ISS isolates

Average nucleotide identity (ANI) analysis revealed that all *K. quasipneumoniae* subsp. *similipneumoniae* strains belonging to ST138 shared an ANI of >99.99% (99.998% ± 0.001), indicating that they belong to the same clone that shows spatio-temporal persistence. Strain IIIF3SW-P1 (ST3234) shared an ANI of 99.148% ± 0.001 with the rest of *K. quasipneumoniae* subsp. *similipneumoniae* strains. *K. pneumoniae* strain F3-2P(2) and *K. aerogenes* strain IIIF7SW-P1 where more genomically distant with ANIs of 93.237% ± 2.346 and 86.079% ± 0.041 when compared to the *K. quasipneumoniae* subsp. *similipneumoniae* strains respectively (**Fig. S1**).

### 2.6. Comparative genomics of K. pneumoniae ST101

The *K. pneumoniae* ST101 is a globally distributed clone with over 696 isolates identified in over 33 countries. *K. pneumoniae* ST101 is widely associated with hypervirulent pathotypes. Overall, the ST101 lineage genomes exhibited remarkable genetic similarity (ANI>99.4%; **Fig. 2A)** and are most associated with Europe and South-East Asia (**Fig. 2B**). F3-2P(2), the ISS strain, is most closely related to ERR985162, a genome isolated from sewage waste water in the UK in 2014, with an ANI of 99.86%. Phylogenetic analysis revealed that F3-2P(2) belongs to a distinct sub-clade within the ST101 lineage, along with 41 other genomes (**Fig. 2C)**. Out of the 696 *K. pneumoniae* ST101 isolates reported worldwide, 87.07% (606/696) were found to carry the *ybt* gene cluster, which is a determinant of virulence associated with the presence of yersiniabactin, suggesting that yersiniabactin plays a key role in the pathogenicity of this clone. Among these isolates, only nine encoded for additional siderophore virulence factors such as colibactin or aerobactin. Moreover, 98.2% (595/606) of the ybt-positive genomes encoded for ybt9, as part of a horizontally acquired ICEKp3. The integrative conjugative element ICEKp5, which was identified in the ISS isolate F3-2P(2) and carries the ybt14 locus, was only found in four other ST101 genomes.

**Figure 2:**
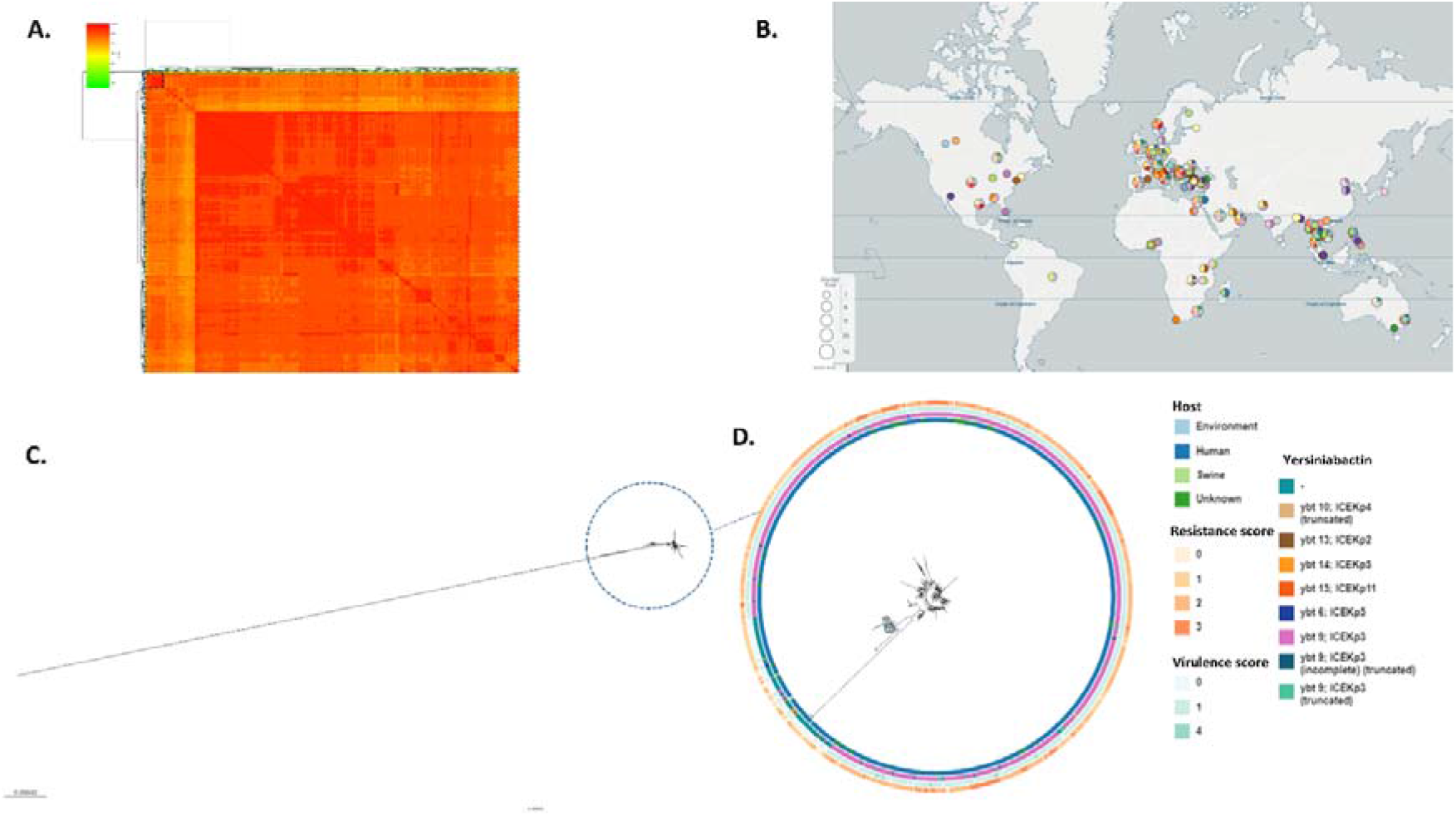
Phylogenetic relationships within *K. pneumoniae* ST101. **A.** Average nucleotide identity (ANI) between the genome of K. pneumoniae ISS strain F3-2P(2), the *K. pneumoniae* typing strain (ATCC13883) and all *K. pneumoniae* genomes of ST 101 (n=696). The sub-clade of F3-2P(2) (n=41) is highlighted **B.** Global distribution of the *K. pneumoniae* ST 101 isolates. **C.** Core-genome phylogenetic relatedness of the *K. pneumoniae* genomes including the typing strain. **D.** Core-genome phylogenetic relatedness of the *K. pneumoniae* genomes excluding the typing strain. The clade within which strain F3-2P(2) is identified is highlighted. An interactive dashboard can be found <u>here</u>.

### 2.7. Comparative genomics of K. aerogenes ST103

The genomes of the *K. aerogenes* ST103 isolates (n=7), including the ISS strain IIIF7SW-P1, were found to be closely related with an ANI >99.8% (**Fig. 3A**). The associated ST103 isolates were spatially restricted, with isolates obtained from human hosts in the USA (n=4) and Japan (n=1) (**Fig. 3B**). The genetically closest strain to IIIF7SW-P1 was SRR5666484, which was isolated from a human with a UTI in the USA in 2015 (ANI=99.97%). Based on core-genome phylogeny, IIIF7SW-P1 formed a distinct node within the ST103 clone along with uropathogenic strain SRR5666484.

**Figure 3:**
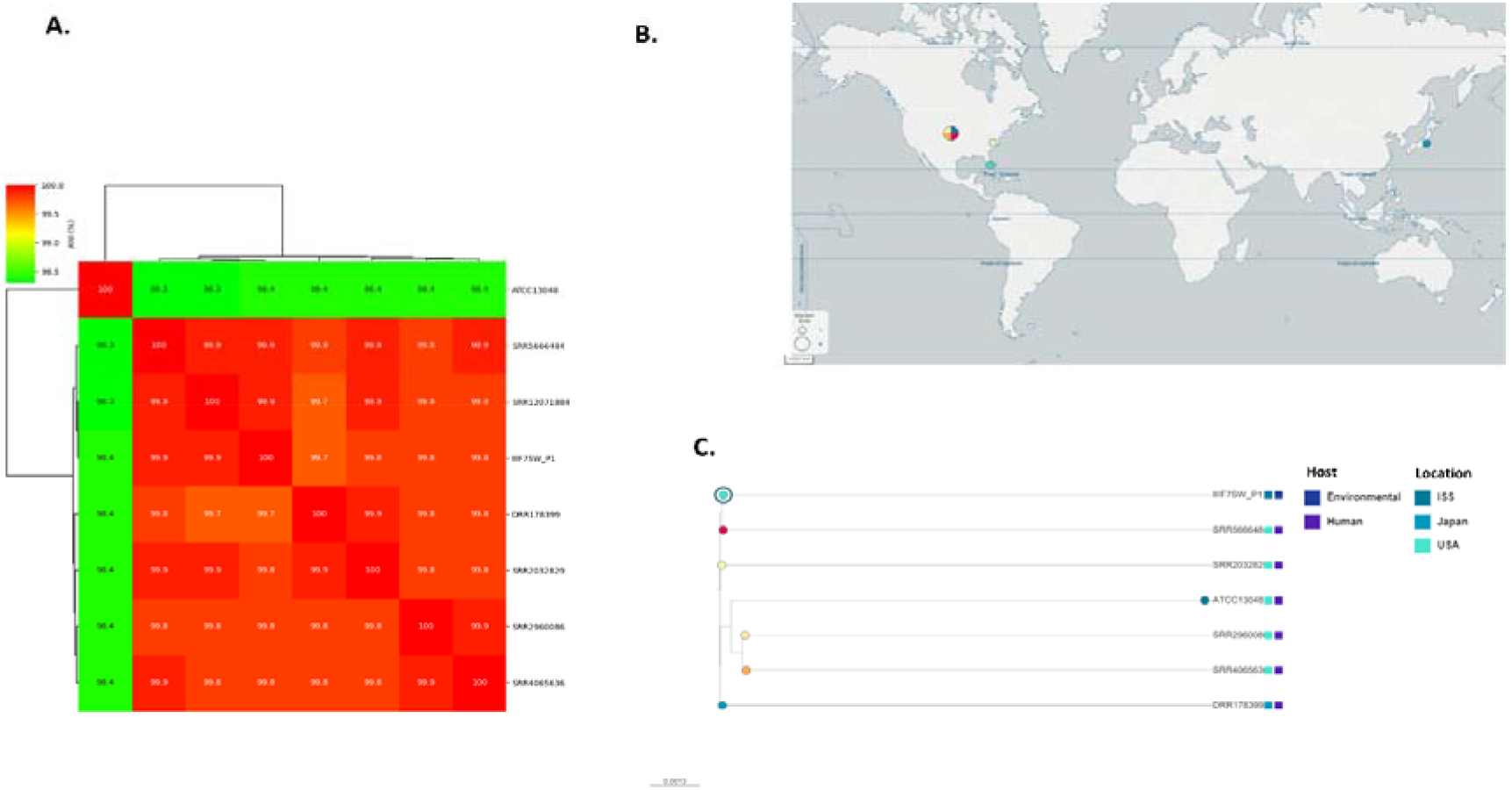
Phylogenetic relationships within *K. aerogenes* ST103. **A.** Average nucleotide identity (ANI) between the genome of *K. aerogenes* ISS strain IIIF7SW-P1, the genome of *K. aerogenes* typing strain ATCC13048 and all K. aerogenes genomes of ST103 (n=6). **B.** Temporal dissemination of the *K. aerogenes* ST103 genomes. **C.** Core-genome phylogenetic relatedness of the *K. aerogenes* ST103 genomes. An interactive dashboard can be found <u>here</u>.

### 2.8. Comparative genomics of K. quasipneumoniae ST3234

*K. quasipneumoniae* ST3234 is a rarely identified clone with only 2 isolates present at the Pathogenwatch database. The genomes of *K. quasipneumoniae* IIIF3SW-P1 and two genomes from isolates from Oxford, UK were found to be highly similar, with an average nucleotide identity (ANI) > 99.8% (**Fig. 4A**). The two UK strains were isolated from cases of invasive bloodstream infections **(Fig. 4B**). When analyzed using core-genome phulogenomics, IIIF3SW-P1 was part of a separate node from the two strains from Oxford (**Fig. 4C**).

**Figure 4:**
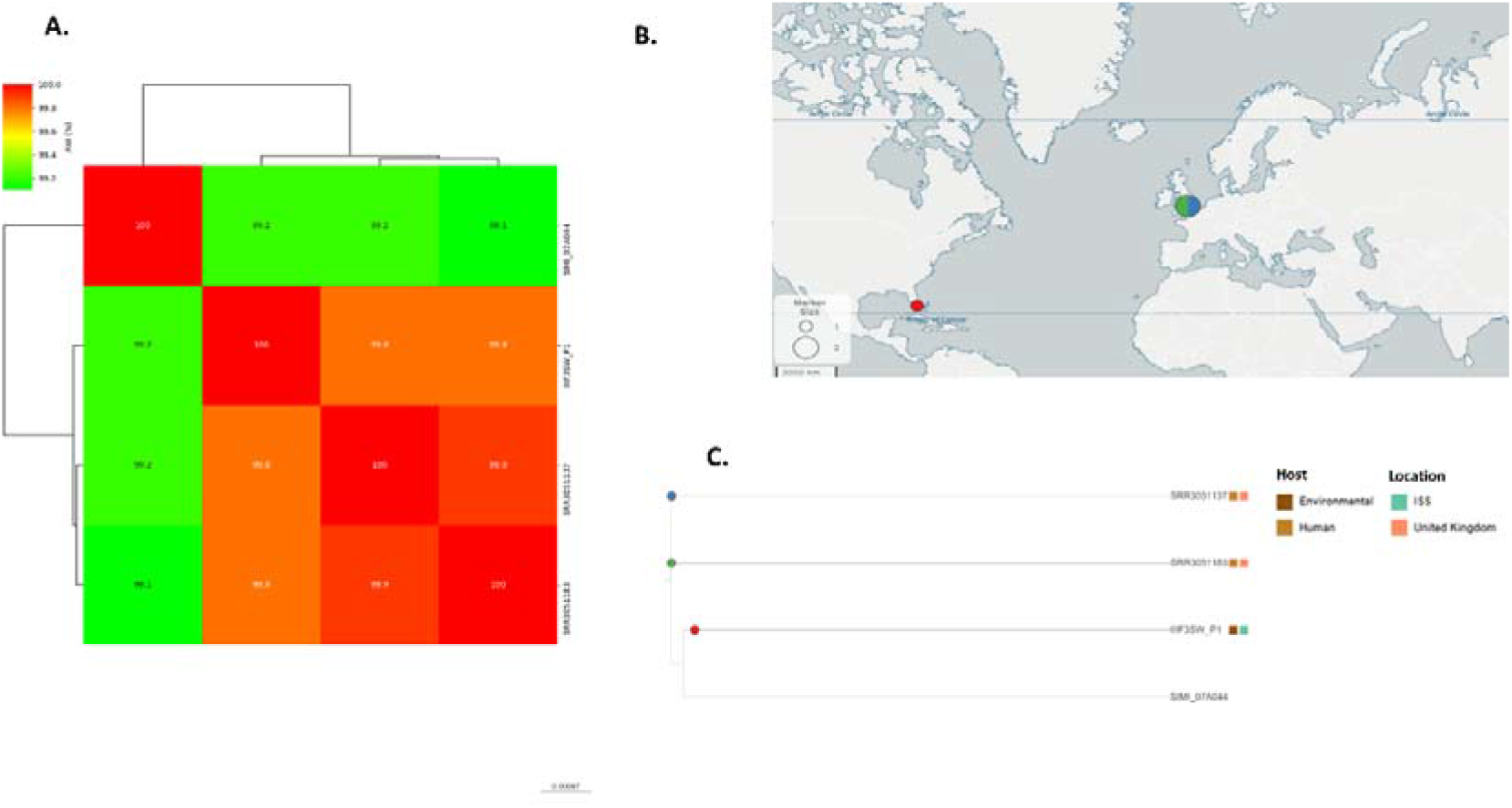
Phylogenetic relationships within *K. quasipneumoniae* ST3234. **A.** Average nucleotide identity (ANI) between the genome of *K. quasipneumoniae* ISS strain IIIIF3SW-P1, the genome of *K. quasipneumoniae* typing strain 07A044 and all *K. quasipneumoniae* genomes of ST3234 (n=2). **B.** Temporal dissemination of the *K. quasipneumoniae* ST3234 genomes. **C.** Core-genome phylogenetic relatedness of the *K. quasipneumoniae* ST3234 genomes. An interactive dashboard can be found <u>here</u>.

### 2.9 Comparative genomics of K. quasipneumoniae ST138

The *K. quasipneumoniae* ST138 lineage showed a wider spatial distribution and higher genetic diversity compared to ST3234, with ANI >99.3% (**Fig. 5A,B**). The closest genome to the seven ISS *K. quasipneumoniae* ST138 isolates, was ERR4635453, isolated from a human host in Tacloban city, Philippines in 2017, with ANI=99.77%. The ISS strains (n=7) formed a distinct clade within the ST138 lineage based on core genome phylogeny, indicating evolutionary divergence from the main ST138 lineage.

**Figure 5:**
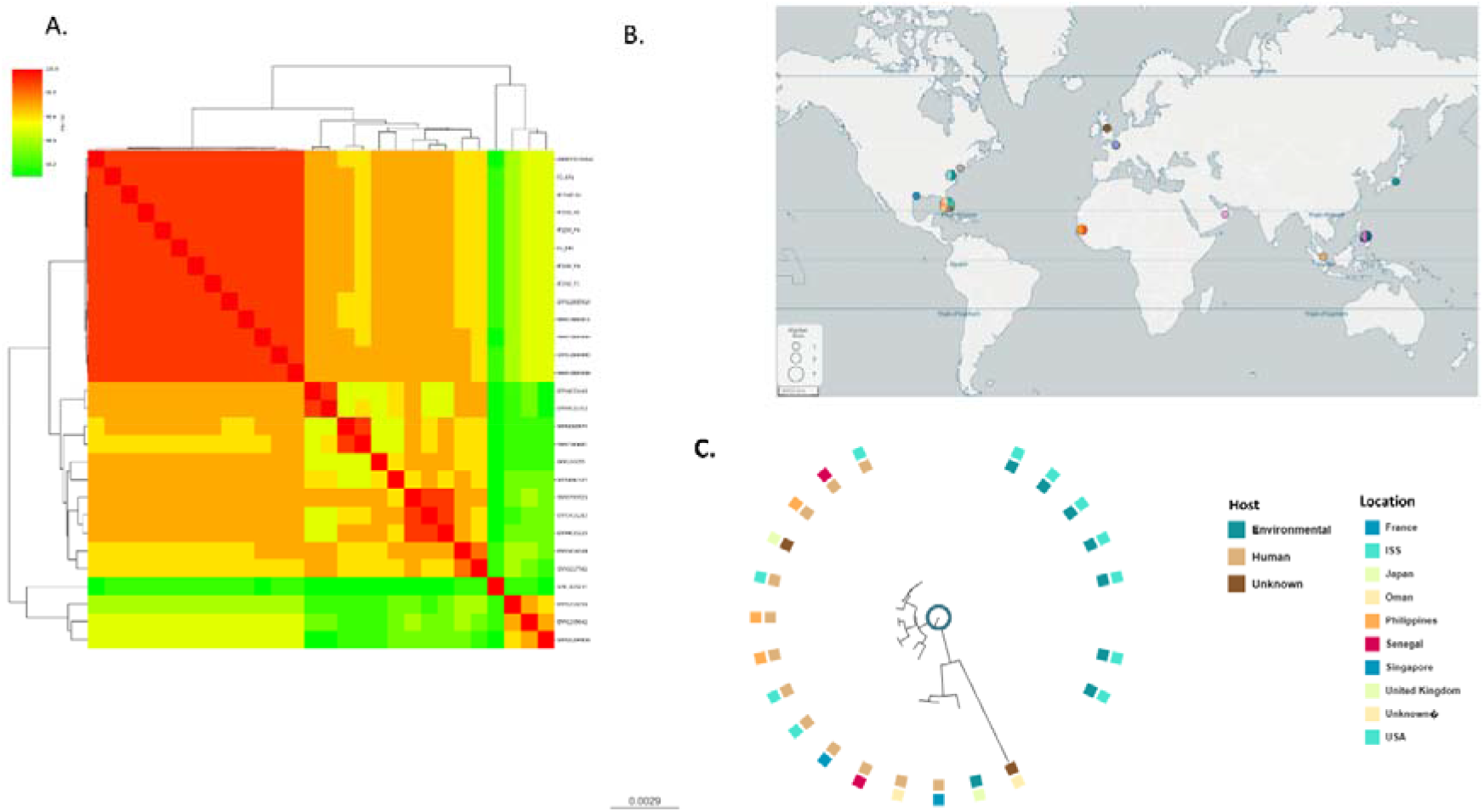
Phylogenetic relationships within K. quasipneumoniae ST138. **A.** Average nucleotide identity (ANI) between the genome of *K. quasipneumoniae* ST138 ISS strains (n=7) the genome of *K. quasipneumoniae* typing strain 07A044 and all *K. quasipneumoniae* genomes of ST138 (n=20). **B.** Temporal dissemination of the *K.* quasipneumoniae ST138 genomes. **C.** Core-genome phylogenetic relatedness of the *K. quasipneumoniae* ST138 genomes. The clade of the ISS isolates is highlighted. An interactive dashboard can be found <u>here</u>.

### 2.10. Antimicrobial resistance genes

The ISS strains (n=10) were found to carry an average of 22.70 ± 0.64 antimicrobial resistance genes (ARGs) each (**Fig. 6A**). Among the genomes, *K. pneumoniae* F3-2P(2) (ST101) was identified to carry the beta-lactamase gene *blaSHV-1*, which is known to confer resistance to penam and some cephalosporin antibiotics (**Fig. 6B**). In the *K. quasipneumoniae* genomes of ST138, chromosomal *blaOKP-B-8* was identified, which confers resistance to first-generation cephems. *K. quasipneumoniae* strain IIIF3SW-P1 (ST3234) was found to carry a *blaOKP-B-3* allelic variant, also conferring resistance to penam and some cephalosporin antibiotics. *K. aerogenes* strain IIIF7SW-P1 (ST103) carried three missense mutations in the outer membrane protein *ompK* gene, which are linked with increased resistance to cephalosporins and would explain the phenotypic resistance to cefoxitin (**Table S5**).

**Figure 6:**
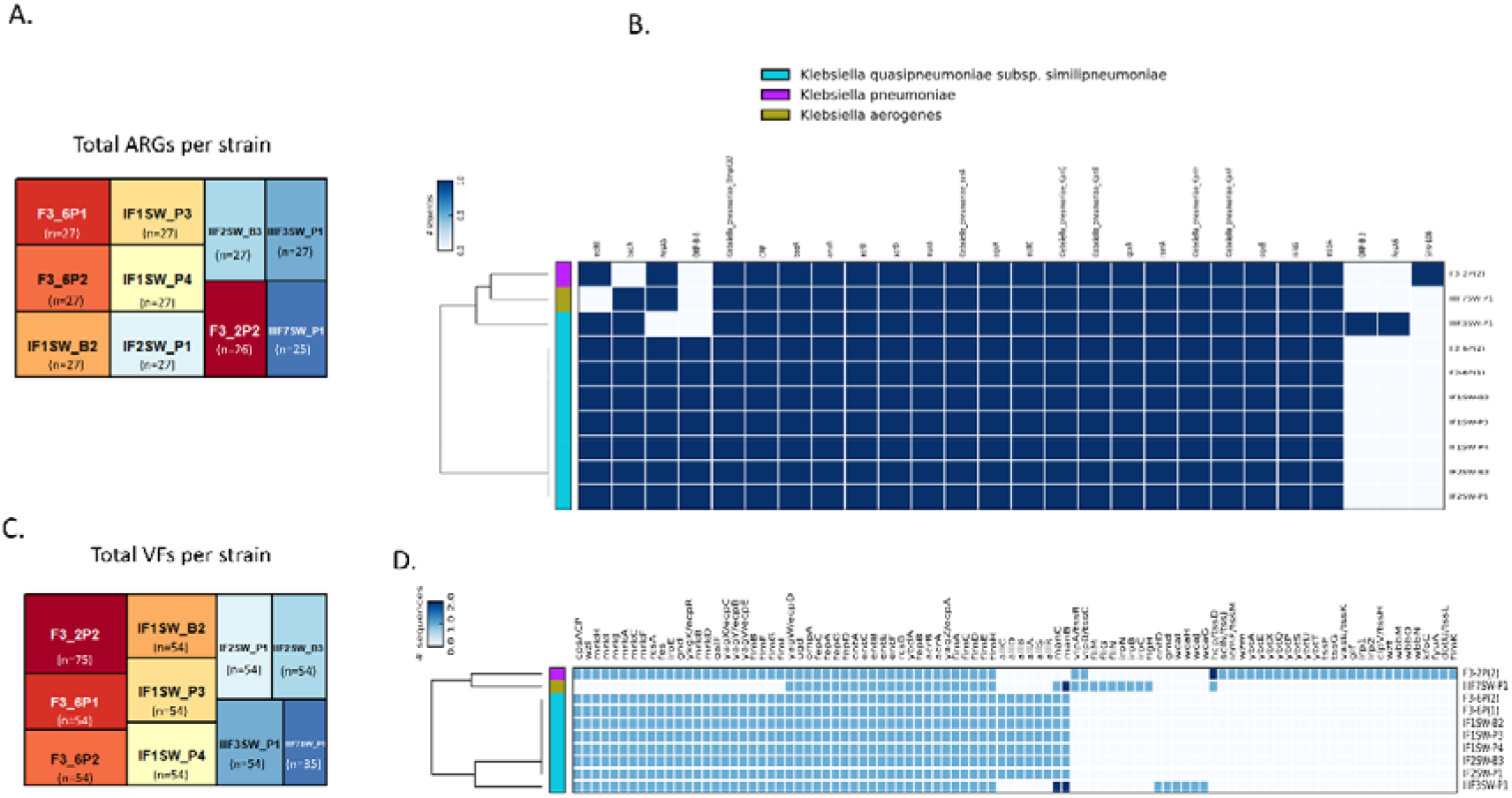
Genetic determinants of resistance and virulence. **A.** Total number of antimicrobial resistance genes (ARGs) identified per ISS originating *Klebsiella* strain. **B.** Heatmap distribution of ARG per isolate. **C.** Total number of virulence factors (VFs) identified per ISS originating *Klebsiella* strain. **D.** Heatmap distribution of VFs per isolate.

### 2.11. Genetic determinants of virulence

The ISS strains (n=10) were found to possess an average of 54.20 ± 8.95 virulence factor (VF) genes each (**Fig. 6C**). *K. pneumoniae* strain F3-2P(2) contained 75 VFs, including the virulence loci for enterobactin and yersiniabactin. The yersiniabactin pathogenicity island (ybt-14 lineage) was part of a self-mobilizable ICE (ICEKp5). The *ybtQ* gene amplicon (234-bp) was also detected using a multiplex PCR assay for the F3-2P(2) strain. Moreover, the iron regulatory protein-encoding genes *irp1* and *irp2* were identified as part of ICEKp5. As a result, strain F3-2P(2) had a virulence score of 1 based on Kleborate’s scoring system [10]. The *wb* gene cluster related to O1 antigen production, commonly found in clinical isolates, was also identified in strain F3-2P(2).

In addition, all 10 ISS *Klebsiella* strains encoded for the virulence-associated enterotoxin (*entA/B/C/E/F*). *K. quasipneumoniae* strain IIIF3SW-P1 (ST3234) encoded the enterotoxin *entD* and (*wcaG/H/I/J*), a gene cluster involved in the biosynthesis of the outer core lipopolysaccharide. *K. aerogenes* strain IIIF7SW-P1 was the only strain identified to contain the *flgH/G/M/N* genes, which encode for regions of the bacterial flagellar switch protein (**Fig. 6D**).

## 3. Discussion

This study highlights the presence of *Klebsiella* isolates associated with human pathogenic lineages and reports on the first identification of a yersiniabactin-encoding *K. pneumoniae* and *Klebsiella* associated plasmids aboard the ISS. Of note, isolate F3-2P(2) belongs to the emerging high-risk clone ST101, which is linked with carbapenem and colistin resistance and has a hypervirulent *K. pneumoniae* (hvKp) pathotype(24),(25). The ST101 clone has been implicated in numerous outbreaks of bloodstream infections worldwide, including in southern Italy and India. (26, 27) ST101 has been identified in isolates from over 33 countries, representing a persistent public health threat. Infections caused by ST101 have been associated with an 11% increase in mortality compared to non-ST101 infections.(28). Widely disseminated, high risk clones like ST101 are known to be able to horizontally acquire hypervirulence gene clusters, potentially leading to the emergence of hypervirulent clonal lineages(29). In accordance with this, our analysis suggests that the predominant virulence factor associated with ST101 is yersiniabactin, as part of a horizontally acquired and self-mobilizable ICEKp.

*K. pneumoniae* strain F3-2P(2) was identified to encode for 75 VF genetic determinants, including the siderophore genes (*irp1/2*), the enterobactin gene cluster and the yersiniabactin (ybt-14) gene cluster as part of a self-mobilizable ICEKp5. *K. pneumoniae* clinical isolates harboring yersiniabactin on ICEKp5 are known to prevail in intensive care units and tertiary hospital settings and are linked with invasive infections(30–32). Among 696 ST101 genomes available on the Pathogenwatch surveillance database, only four carried the same ICEKp5 harboring the ybt-14 gene cluster. These strains were distributed widely in different countries such as USA, China, and Philippines. The mode of transportation of a potential pathogen such as *K. pneumoniae* F3-2P(2) to the ISS could be through cargo or crew, although relevant crew data were not available as the National Aeronautics and Space Administration (NASA) does not mandate the monitoring of microbial diversity.

The genome of *K. pneumoniae* strain F3-2P(2) also revealed the presence of an IncFIIK conjugative plasmid containing several heavy-metal resistance and thermoresistance genes. Although ESBL CTX-M-15 encoding plasmids are commonly associated with the FIIK-9 replicon, the plasmid of F3-2P(2) lacked ESBLs likely due to the absence of antibiotic-induced selective pressure(33). This strain was isolated from the waste and hygiene compartment (WHC) of the ISS. The role of the WHC is the removal and containment of human solid waste and urine. While it is unclear if the strain originated from the ISS crew or cargo, *K. pneumoniae* strains have been reported to survive on surfaces for up to 6 weeks, with the longest survival rates on stainless steel (34),(35). Human infection with hvKp isolates with both enterobactin and yersiniabactin virulence loci present, could result to multi-site infection due to metastatic spread (36). Furthermore, the hypervirulent, multimetal resistance and thermotolerant genomic profile of the *K. pneumoniae* F3-2P(2) strain, could possibly allow it to evade the existing disinfecting protocols aboard the ISS. The presence of the resistance and hypervirulent determinants in MGEs further poses a consideration for their dissemination across the ISS microbiome.

The dominance of *K. quasipneumoniae* ST138 on the ISS environment was also evidenced by the isolation of seven strains over 18 months from three different locations (Permanent Multipurpose Module, Cupola, and WHC). Despite the spatial and temporal distribution of their collection, the ST138 isolates showed remarkable genetic similarity, suggesting the possible propagation of a single strain in the closed system for a prolonged period. The ST138 isolates shared several chromosomal genetic determinants of resistance to disinfectants and heavy metals, including resistance to quaternary ammonium compounds (QACs), peroxide, fosmidomycin, methyl viologen, tellurite, nickel, cobalt, zinc and the multiple stress resistance protein BhsA encoded by *bhsA*. These determinants may confer adaptability advantages to the ST138 lineage during spaceflight, given most cleaning agents and disinfectants contain several heavy metals and QACs(37). On Earth *K. quasipnuemoniae* ST138 strains have been isolated from untreated and treated water in a wastewater treatment plant in Slovenia(38). ESBL cefotaximases (CTXM-9) producing isolates of *K. quasipneumoniae* ST138 have also been reported to cause community onset infections in China(38). Overall. *K. quasipnuemoniae* ST138, shows temporal and spatial persistence aboard the ISS, more than any other *Klebsiella* lineage. The ISS *K. quasipnuemoniae* ST138 strains lacked any MGEs and therefore their adaptability advantage could be intrinsic. Further research is required to elucidate the properties of this lineage, which allow it to adapt and propagate to the high-pressure environment of the ISS.

*K. quasipneumoniae* subsp. *similipneumoniae* strain IIIF3SW-P1 was the only ST3234 strain identified on ISS, and it was isolated from the crew exercise platform (ARED). There is not much known about the dissemination and persistence of *K. quasipneumoniae* ST3234 as only two isolates have been reported on surveillance databases. Both Earthly strains were associated with invasive blood stream infections in humans and were isolated at John Radcliffe Hospital in Oxford, UK in between 2009 and 2012.

*K. aerogenes* strain IIIF7SW-P1 (ST103) was isolated from the ISS Lab3 in 2015. This strain belongs to the globally rare *K. aerogenes* ST103 lineage, with only five strains deposited in surveillance databases, all of which are associated with UTIs and reported in the USA. Interestingly, IIIF7SW-P1 harbored two uncharacterized plasmids, one of which (pIIF7SW-P1_1) was predicted to be mobilizable and contained the pilL gene responsible for thin pilus biogenesis(39). PilL is known to promote bacterial adhesion and colonization, providing a selective advantage in the high selective pressure environment of the ISS. The second plasmid (pIIF7SW-P1_2) was a high copy plasmid, harboring the heat shock gene htrC, which has been linked with increased thermoresistance in *E. coli*(40). The genetic makeup of IIIF7SW-P1 suggests that it may have acquired plasmids enabling it to survive and persist in the challenging conditions of the ISS, which warrants further investigation.

In this study, the first complete genomic characterization of *Klebsiella* strains isolated from the ISS was conducted. This study reports on the genomic makeup of ten *Klebsiella* isolates, revealing an abundance of heavy metal and QAC resistance genes amongst the genomes, similar to the ones seen in clinical settings where the use of disinfectants is intense(37). However, clinically relevant ESBLs and carbapenemase-producing genes were notably absent. This study reports for the first time on the isolation of yersiniabactin encoding *K. pneumoniae* from a human associated pathogenic lineage aboard the ISS as well as on the persistence and possible genomic divergence of *K. quasipneumoniae* ST138 isolates. The prevalence and seemingly adaptability of *K. quasipneumoniae* ST138 in spaceflight conditions should be considered for further investigation. Additional epidemiological studies are needed to map and understand the prevalent *Klebsiella* lineages that could persist during prolonged spaceflight. Ongoing surveillance of potentially problematic STs is feasible on the ISS due to its small crew size and could help mitigate associated risks. These efforts will aid in developing suitable countermeasures for eradicating potentially problematic pathogens in closed habitats for future human missions to the Moon, Mars, and beyond.

## 4. Materials & Methods

### 4.1. Isolation of the strains from the ISS

Various locations were sampled on the ISS using polyester wipes, and the metadata associated with the samples and their collections were published elsewhere(41). Briefly, each wipe was aseptically removed from the zip lock bag and transferred to a 500 mL bottle containing 200 mL of sterile phosphate-buffered saline (PBS; pH 7.4) and concentrated with a Concentrating Pipette (Innova Prep, Drexel, MO) using a 0.22 µm Hollow Fiber Polysulfone tips (Cat #: CC08022). Suitable aliquots (100 µL) of each sample were plated on Reasoner’s 2A agar in duplicate and incubated at 25 °C for seven days, and well-matured colonies were picked, archived in semisolid R2A (agar media diluted 1:10) and stored at room temperature. A loopful of purified microbial culture was directly subjected to PCR. The targeted fragment was amplified (colony PCR) to amplify the 1.5 kb 16S rRNA gene to identify bacterial strains. The following primers were used for the 16S rRNA gene amplification: the forward primer, 27F (5’-AGA GTT TGA TCC TGG CTC AG-3’), and the reverse primer, 1492R (5’-GGT TAC CTT GTT ACG ACT T-3’)(42, 43). The PCR conditions were as follows: denaturation at 95 °C for 5 min, followed by 35 cycles consisting of denaturation at 95 °C for 50 s, annealing at 55 °C for 50 s, and extension at 72 °C for 1 min 30 s and finalized by extension at 72 °C for 10 min. The sequencing was performed using 27F, and 1492R primers and the sequences were assembled using SeqMan Pro from DNAStar Lasergene Package (DNASTAR Inc., Madison, WI). The bacterial sequences were searched against EzTaxon-e database(44) and the initial identification was based on the closest percentage similarity (>97%) to previously identified microbial type strains.

### 4.2. DNA extraction and sequencing

Cultures of the 10 *Klebsiella* strains were grown overnight on MacConkey agar plates at 37°C. DNA extraction was performed using the QIAamp DNA Mini Kit (Qiagen, Germany) according to the manufacturer’s instructions, with no pre-enrichment steps. The extracted DNA was quantified using a Qubit fluorometer (Thermo-Fisher Scientific, USA) and the dsDNA High Sensitivity kit (Thermo-Fisher Scientific, USA). DNA purity was evaluated by measuring the absorbance ratio at 260 nm and 280 nm (A260/280) on a Jenway™ Genova Nano Micro-volume Spectrophotometer (ThermoFisher Scientific, USA). For sequencing, DNA samples with a suitable quantity (>12.5 ng/μL) and purity (A260/280 of 1.80-2.00) were selected. Illumina NovaSeq 6000 sequencing was performed on these samples at Oxford’s Genomics Centre (PE150). In addition, long-read sequencing for the DNA of all ten strains was carried out using an Oxford MinIONMk1C platform (Oxford, UK).

### 4.3. Antimicrobial susceptibility testing (AST)

Antimicrobial susceptibilities for the ten *Klebsiella spp*. isolates were determined by the Kirby-Bauer disc diffusion method for a range of 8 β-lactam and 10 non-β-lactam antibiotics as previously described(45) (**Table S6**). The diffusion test was performed on Mueller Hinton (MH) agar as culture medium. Classification into susceptibility categories was determined using the clinical breakpoints provided for Enterobacterales by the Clinical and Laboratory Standards Institute (CLSI) (30^th^ edition)(46).

### 4.4. Bioinformatic analysis

#### Filtering and assembly

For short reads, quality filtering and adapter trimming was carried out using fastp (v0.23.2)(47). Only reads with a Q score of over 20 were retained. For long reads, adapter trimming was conducted using porechop (v0.2.4)(48). Trimmed long reads were quality filtered using <u>filtlong</u> (v0.2.1) and only reads of 1kbp or more were retained. Hybrid genome assembly was conducted using the hybrid assembly pipeline Unicycler (v0.5.0)(49) with default settings. The quality of hybrid assemblies was assessed using QUAST (v5.2.0)(50). Closed genome assemblies were visualized using Bandage (v0.8.1)(51), separating chromosomal and plasmid sequences. Circular visualization of the chromosomes and plasmids coupled including COG classification of their genetic features was conducted using <u>COGclassifier</u> (v1.0.4) and <u>MGCplotter</u> (v1.0.1).

#### Genomic characterisation of resistance, virulence, and plasmids

Genome comparison between the ten ISS strains was also conducted using MGCplotter utilizing *K. quasipneumoniae* strain IF1SW-B2 as a reference genome (v1.0.1) (**Fig. 1**). The mobilization and conjugation potential of the plasmids was predicted using the Mobtyper tool from the MOB-suite software package (v3.1.0)(52). Plasmid ST was determined using the <u>plasmidMLST</u> (v2.0) tool.

The average nucleotide identity (ANI) was determined using <u>ANIclustermap</u> (v1.1.0). The bacterial chromosomes and plasmids identified were scanned for the presence of ARGs and virulence factors (VFs) using <u>Abricate</u> (v.1.0.0) against the comprehensive antibiotic resistance database (CARD)(53) and the virulence factor database (vfdb)(54) respectively. Only results with coverage and identity >90% were considered. Species identification, sequence typing, K and O locus typing, and quantification of the respective virulence and resistance scores were conducted using Kleborate (v2.2.0) (55) and Kaptive (v2.0.3) (56).

#### Large scale comparative genomics

To compare the ISS *Klebsiella* genomes with earthly analogues of the same ST, all available genomes from the STs identified for the ISS strains were retrieved from the Pathogenwatch database (**Table S6**). Their assemblies were annotated using PROKKA (v1.14.5)(57). A pan-genome analysis for each ST, which included the corresponding ISS strains and the species typing strain was conducted using roary (v3.11.2) (58). Core genome alignment was determined using mafft (v7.47.1)(59), and an approximately-maximum-likelihood phylogenetic tree utilizing the generalized time-reversible (GTR) model was constructed using FastTree (60)(v2.1.11). The presence-absence results from the pan-genome analysis and the phylogenetic tree generated were visualised using microreact(61). Data were also visualized using the R programming language.

Dedicated interactive dashboards per ST have been generated and are accessible in the following links: *<u>K. quasipneumoniae ST 3234; K. quasipneumoniae ST138; K. pneumoniae ST101, K. aerogenes ST103</u>*

### 4.5. Hypervirulence multiplex PCR assay

An in-house developed and validated multiplex PCR assay, designed to detect hypervirulence in clinical *K. pneumoniae* strains was used to verify the presence of yersiniabactin identified in strain F3-2P(2). The assay targets the *ybtQ* gene of the yersiniabactin gene cluster, the *iucA* gene of the aerobactin gene cluster, and the *16SrRNA* gene as a positive control (**Table S7**). The multiplex PCR was conducted on a SensQuest Thermocycler with the following conditions: 2mins-50°C, 10mins-95°C; (15secs-95°C, 1min-60°C) x35.

## Data availability

All genomic data presented herein are publicly available in GenBank under BioProject numbers PRJNA635942, PRJNA640688 and PRJNA635942.

## Acknowledgments

Part of the research described in this manuscript was performed at the Jet Propulsion Laboratory, California Institute of Technology, under a contract with NASA. We would like to thank Aleksandra Checinska-Sielaff for isolating the strains. We thank astronauts Captain Terry Virts for collecting samples aboard the ISS and the Implementation Team at NASA Ames Research Center (Fathi Karouia) for coordinating this effort. We also acknowledge the Jet Propulsion Laboratory supercomputing facility staff, notably Narendra J. (Jimmy) Patel and Edward Villanueva, for their continuous support in providing the best possible infrastructure for BIG-DATA analyses. ©2023 California Institute of Technology. Government sponsorship acknowledged.

## Ethics

No ethical approval or consent was needed for this manuscript.

## 16. Author contributions

K.V., G.M., and N.K.S conceived and designed the experiments. A.T. designed and conducted the multiplex PCR assay. L.O’C. and G.M. generated genomic libraries and conducted the long-read sequencing. G.M. and F.MD. conducted the microbiological phenotypic characterisation. G.M., K.V., N.K.S. and F.MD. conducted the genomic analysis. G.M. generated the figures and the manuscript was compiled by all authors. The authors have read and agreed to the published version of the manuscript.

