## Supplementary material for "Genomic analysis reveals the presence of emerging pathogenic *Klebsiella* lineages aboard the International Space Station"

**Table S1:** Antimicrobial Susceptibility Testing (AST) of ISS *Klebsiella* isolates including their ST profile and the isolation site aboard the ISS.

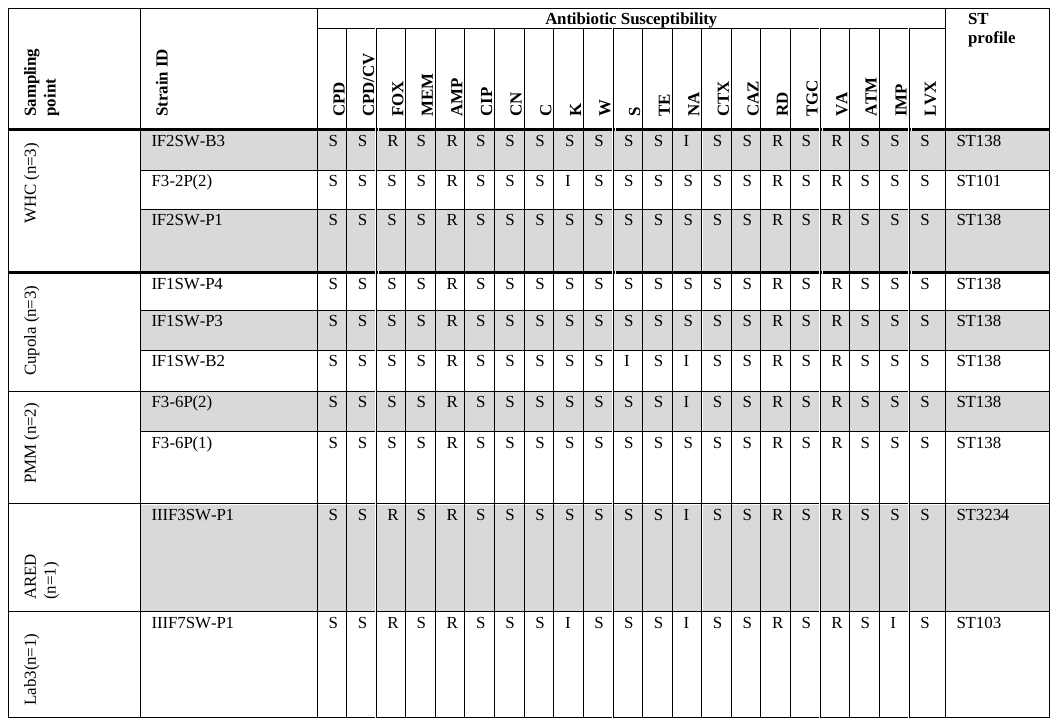

CPD= cefpodoxime, CPD/CV= cefpodoxime/clavulanic acid, FOX = cefoxitin, MEM = meropenem, AMP = ampicillin, CIP = ciprofloxacin, CN = gentamicin, K= kanamycin, W= trimethoprim, S= streptomycin, TE = tetracycline, NA= nalidixic acid, CTX= cefotaxime, CAZ= ceftazidime, RD= rifampicin, TGC= tigecycline, ATM= aztreonam, IMP= imipenem, LVX= levofloxacin, VA= vancomycin, ATM= aztreonam

**Table S2**: Genomic characteristics of the completed genomes: strain name, genome size, sequencing depth, and GC content as well as GenBank accession number.

| **Strain name** | **Genome size (bp)** | **Depth (x)** | **GC content (%)** | **BioSample accession number** | **GenBank accession number(s)** |
| --- | --- | --- | --- | --- | --- |
| F3-6P(2) | 5,234,651 | 745 | 58.07 | SAMN15329710 | CP118276 |
| IF2SW-B3 | 5,234,630 | 914 | 58.07 | SAMN15061189 | CP118271 |
| F3-2P(2*) | 5,311,445 | 514 | 57.22 | SAMN15329758 | CP118405-CP118406 |
| IF1SW-B2 | 5,234,636 | 745 | 58.07 | SAMN15061188 | CP118277 |
| IIIF7SW-P1 | 5,162,861 | 592 | 55.02 | SAMN15344675 | CP118332-CP118334 |
| F3-6P(1) | 5,234,651 | 530 | 58.07 | SAMN15329711 | CP118273 |
| IIIF3SW-P1 | 5,189,883 | 531 | 58.10 | SAMN15329712 | CP118278 |
| IF2SW-P1 | 5,234,636 | 709 | 58.07 | SAMN15061190 | CP118272 |
| IF1SW-P4 | 5,234,636 | 514 | 58.07 | SAMN15061191 | CP118275 |
| IF1SW-P3 | 5,234,636 | 784 | 58.07 | SAMN15061192 | CP118274 |

**Table S3:** Typing of the *Klebsiella* strains included in this study. Their ST, K, and O locus and wzi allelic variants are presented.

| Strain name | Species | ST | K locus | O locus | wzi |
| --- | --- | --- | --- | --- | --- |
| F3-6P(2) | *K.quasipneumoniae* subsp*. Similipneumoniae* | ST138 | K48 | O5 | wzi291 |
| IF2SW-B3 | *K.quasipneumoniae* subsp*. similipneumoniae* | ST138 | K48 | O5 | wzi291 |
| F3-2P(2*) | *K. pneumoniae* | ST101 | unknown (KL106) | O1 | wzi29 |
| IF1SW-B2 | *K.quasipneumoniae* subsp*. similipneumoniae* | ST138 | K48 | O5 | wzi291 |
| IIIF7SW-P1 | *K. aerogenes* | ST103 | unknown (KL119) | O3/O3a | - |
| F3-6P(1) | *K. quasipneumoniae* subsp*. similipneumoniae* | ST138 | K48 | O5 | wzi291 |
| IIIF3SW-P1 | *K. quasipneumoniae* subsp*. similipneumoniae* | ST3234 | K16 | O3/O3a | - |
| IF2SW-P1 | *K. quasipneumoniae* subsp*. similipneumoniae* | ST138 | K48 | O5 | wzi291 |
| IF1SW-P4 | *K. quasipneumoniae* subsp*. similipneumoniae* | ST138 | K48 | O5 | wzi291 |
| IF1SW-P3 | *K. quasipneumoniae* subsp*. similipneumoniae* | ST138 | K48 | O5 | wzi291 |

**Table S4**: Plasmid characterisation including their replicon, relaxase and mpf types as well as predictions for their mobility potential.

| **Plasmid ID** | **Replicon type** | **Relaxase type** | **mpf type** | **Mobility** |
| --- | --- | --- | --- | --- |
| pF3-2P2 | IncFIIK | MOB-F | MPF-F | Conjugative |
| pIIF7SW-P1_1 | Rep_cluster 1367 | MOB-C | - | Mobilisable |
| pIIF7SW-P1_1 | Unknown | - | - | Non-mobilisable |

**Table S5**: Missense mutations associated with cephalosporin resistance in the ompK gene of *K.aerogenes* strain IIIF7SW-P1.

| **Mutation** | **Nucleotide change** | **Amino acid change** |
| --- | --- | --- |
| ompK36 p.N49S | AAC->AGC | N->S |
| ompK36 p.A36S | GCA->TCC | A->S |
| ompK36 p.T184P | ACC->CCA | T->P |

**Table S6**: *Klebsiella* genomes available on Pathognewatch database for each of the ST identified aboard the ISS. The lineage dissemination of each ST is also presented.

| **Species** | **ST** | **Pathogenwatch database genomes available** | **Lineage dissemination (countries where the ST has been identified)** |
| --- | --- | --- | --- |
| *K. pneumoniae* | 101 | 696 | 33 |
| *K. quasipneumoniae* | 138 | 20 | 8 |
| *K. quasipneumoniae* | 3234 | 2 | 1 |
| *K. aerogenes* | 103 | 6 | 2 |

**Table S7**: hvKp multiplex PCR primes and probes: Gene targets for aerobactin (*iucA*) and yersiniabactin (*ybtQ*). Probes are dual labelled and contain a reported dye FAM/JOE (5’ end) and a quencher BHQ1 (3’ end).

| **Target Gene** | **Forward Primer** | **Reverse Primer** | **Probe** |
| --- | --- | --- | --- |
| ***iucA*** | TTTCTCCCCAACCCAGCATC | CGCCAAAGTTCAACGCTTTTTC | 5’-[FAM] AGC CAC GCC CAT ACG CAG CAG GCA [BHQ1]-3’ |
| ***ybtQ*** | ATGTTCTCCAGATGAAAAGCC | CCAATACTGAATCGCTGTCC | 5’-[JOE] CGA ATG TCA CGC AGG CTC CAC GCC AGA A [BHQ1]-3’ |
| ***16SrDNA*** | TGGAGCATGTGGTTTAATTCGA | TGCGGGACTTAACCCAACA | 5’-[JOE] CACGAGCTGACGACARCCATGCA [BHQ1]-3’ |

**Figure S1**: Average nucleotide identity (ANI) analysis of ISS strains. ANI analysis of *Klebsiella* strains isolated aboard the ISS (n=10).
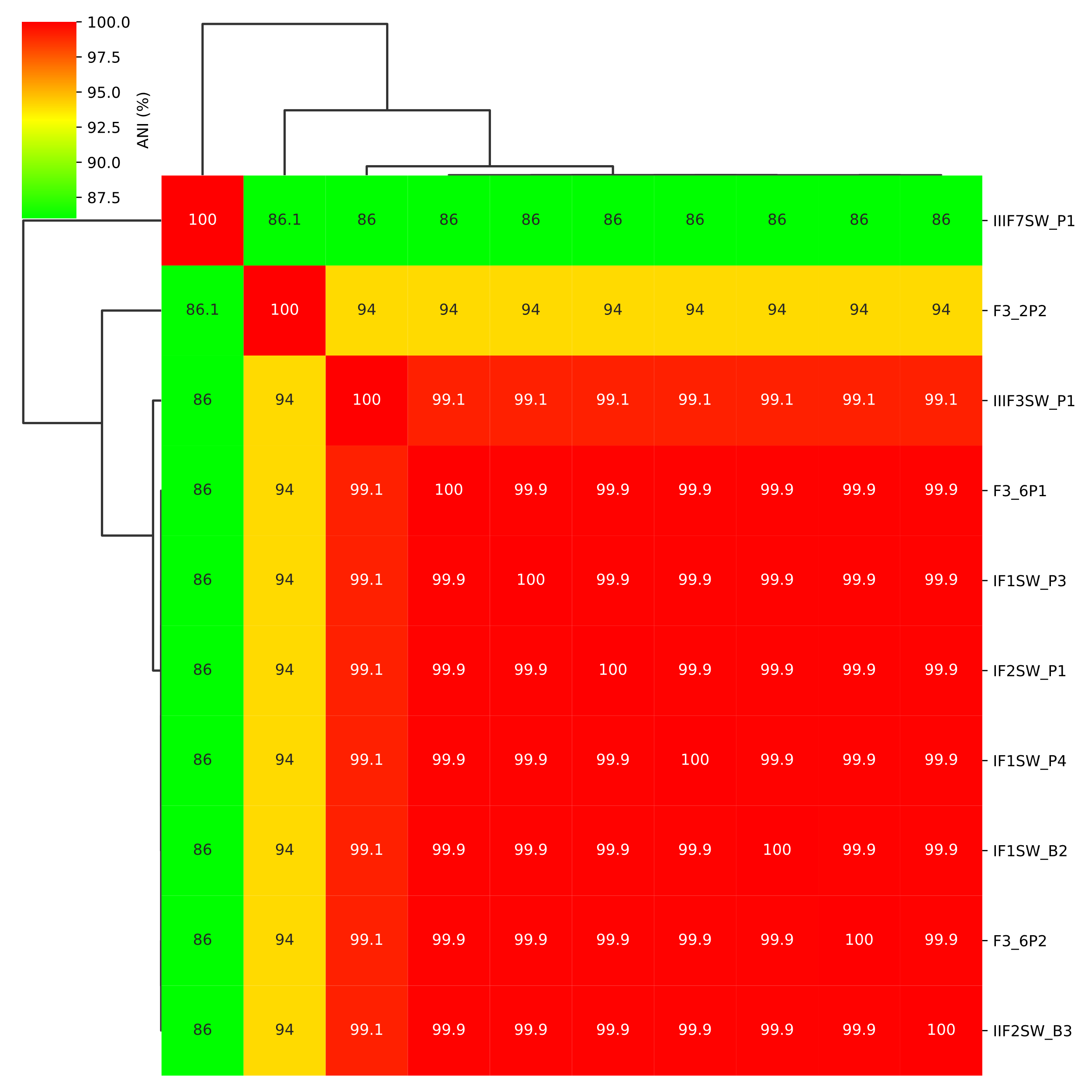
